# Is Democracy the Right System? Collaborative Approaches to Building an Engaged RDM Community

**DOI:** 10.1101/103895

**Authors:** Marta Teperek, Rosie Higman, Danny Kingsley

**Affiliations:** Office of Scholarly Communication, University of Cambridge

## Abstract

When developing new products, tools or services, one always need to think about the end users to ensure a wide-spread adoption. While this applies equally to services developed at higher education institutions, sometimes these services are driven by policies and not by needs of end users. This policy-driven approach can prove challenging for building effective community engagement. The initial development of Research Data Management support services at the University of Cambridge was policy-driven and subsequently failed in the first instance to engage the community of researchers for whom these services were created.

In this practice paper we will describe the initial approach undertaken at Cambridge when developing RDM services, the results of this approach and lessons learnt. We will then provide an overview of alternative, democratic strategies employed and their positive effects on community engagement. We will summarise by performing a cost-benefit analysis of the two approaches. This paper might be a useful case study for any institutions aiming to develop central support services for researchers, with conclusions applicable to the wide sector, and extending beyond Research Data Management services.

## Introduction

When innovators develop new products, the first questions they ask are about what problems the product is intended to solve. This is to achieve the end goal – widespread adoption as the measure of successful innovation. In other words, looking at complex issues from the perspective of end users and their problems is at the core of entrepreneurship (Ries, 2011). Interestingly however, library environments have in the past tended to maintain the status quo and avoided changes and innovation unless necessary (White, 1987). Therefore, sometimes external policies become drivers for change in libraries. In many libraries, and in higher education institutions overall, the recent trend for funders’ policies requiring researchers to manage and share their research data were the drivers for development of support services for research data management (RDM) (Cox & Pinfield, 2014; Dietrich, Adamus, Miner, & Steinhart, 2012; Jones, Pryor, & Whyte, 2013a). A similar policy-driven approach to RDM services development was applied at the University of Cambridge (Open Access Project Board, 2014). However, the initial top-down approach based on meeting policy requirements, without supporting users with appropriate resources (Pryor, 2012) and without trying to break down and understand the complex issues of research data management (Awre et al., 2015), failed to engage researchers at the University of Cambridge. Learning from that experience, several more democratic and end-user focused strategies were implemented instead. These turned out to be much more successful in building researcher community engagement. However, they also proved to be substantially more resource-intensive.

In this paper we will describe and compare the two different approaches towards RDM service development which were used at the University of Cambridge: the policy-driven, top-down approach and more democratic, bottom-up strategies. We will reflect on advantages and disadvantages of both solutions and we will make recommendations about the use of these strategies in development of library services, extending beyond mere RDM support.

## Unsuccessful top-down, policy-driven approach

The initial creation of the Research Data Management Policy Framework the University of Cambridge was largely driven by expectations about data management and sharing from the Engineering and Physical Sciences Research Council (EPSRC) (EPSRC, 2014; Open Access Project Board, 2014; University of Cambridge, 2015). Many other institutions in the United Kingdom adopted a similar policy-driven approach (Weigert, Jones, Duke, & Rans, 2015). The EPSRC requires that all papers acknowledging its funding have a clear statement on data accessibility and that research institutions provide adequate infrastructure to support researchers in effectively managing and sharing their research data. Additionally, researchers and institutions which fail to comply with the EPSRC policy, face potential sanctions from the funder (Ryan, 2015). Therefore, there were several reasons for the initial adoption of a top-down, policy-driven approach at Cambridge:

- The EPSRC is one of the major funders at the University of Cambridge and not complying with the funder’s policy meant that the University might suffer from a substantial income loss – a top-down approach was needed to ensure that both senior University management, as well as researchers, recognised the risk.
- Support services for RDM at Cambridge were underdeveloped (Pryor, 2012). Therefore, the top-down approach and endorsement from senior management offered the possibility of a quick development and roll-out of required services.
- The University of Cambridge is a large, research-intensive institution, with a complex organisational structure of schools, departments and colleges (University of Cambridge, 2017). Hence, a simple, top-down arrangement presented an attractive opportunity for a potential fast and effective message delivery to all researchers and research staff.

We started by organising a series of information sessions, to which we invited researchers, research staff and students. The main message delivered at these sessions was that research data needs to be shared due to funders’ requirements. However, we did not explain to researchers why they should adhere to funders’ requirements, why these policies were introduced by funders in the first place and what the problems these RDM policies were trying to solve (Teperek & Kingsley, 2015b). Additionally, our initial approach was not accompanied by new resources or new services developed and we also did not consult researchers on their experience and views on data management and sharing.

Our initial presentations were perceived by the researcher community as yet a new requirement or another ‘checkbox’ activity, dictated by funders and by the central University administration. Without understanding *why* these policies were introduced, *what problems* they were trying to solve and without appropriate tools to help researchers improve their data management and sharing practice, researchers were disinclined to invest their time and effort in research data management and sharing. We needed to change our approach in order to engage the community and to avoid the risk of developing policies, which will never be practically implemented.

## Efforts to better understand the research community

In order to change our approach and to better tailor it to researchers’ needs, we have invested considerable time and effort in trying to better understand the current practice of research data management in Cambridge and the gaps in RDM support which would need to be filled in order to enable our research community to effectively manage and share their research data. We used several approaches to achieve this:

- Direct discussions with researchers
- Structured interviews and surveys
- Open door meetings with funders

### Direct discussions with researchers

Since January 2015 the team have spoken with over 2,000 researchers across the University, during seventy five separate discussion sessions about research data management. Some of these sessions were held centrally, but most were organised at individual departments (visiting researchers where they work and where they create data). At least two team members would attend these sessions. One would be responsible for giving a short introduction to data management and sharing and for facilitating discussions with researchers, whereas the other team member would note all the questions received. This systematic approach allowed us to create a database of Frequently Asked Questions (Teperek & Kingsley, 2015a). The answers provided in this database were subsequently checked by several funding bodies to ensure the correct information was being conveyed. This was beneficial in several ways. First, the list of FAQs proved to be an effective resource for researchers allowing them to quickly find answers to questions without the need of emailing or calling the support staff. Second, researchers who saw that their questions were not dismissed, but that instead they were answered, recorded and used as a resource for their peers, started to see the benefit of engaging with the service development. Third, asking funders to review the answers not only provided additional credibility to FAQs, but also helped building effective engagement with funding bodies, who were in turn also interested to learn what questions researchers had about their policies. Finally, this approach allowed us to understand the barriers to, and motivations for, good data management and sharing practices.

### Structured interviews and surveys

We followed recommendations developed by the Digital Curation Centre suggesting that the community should shape RDM services (Jones, Pryor, & Whyte, 2013b). We conducted a series of structured interviews and surveys on how the RDM services should look (Johnson, Chiarelli, & Parsons, 2016; Teperek, 2015a). Importantly, each time we asked questions, we explained to researchers why we were asking these questions and our plans to act on the feedback received. Knowing that responses will shape RDM services provided a motivation for the future end users to take part in these surveys. Survey results indicated that the top research data management needs among our research community were: an easily accessible, central information on RDM, training and support in data management across the whole research lifecycle and an easy to use data repository to share research data.

### Open door meetings with funders

To further understand researchers’ problems with research data management and sharing and to ensure that they receive sufficient consideration, we also organised several open door meetings where we allowed researchers to ask questions about data management and sharing directly of funders. Some of the major University funders were invited to these meetings: the Engineering and Physical Sciences Research Council, the Biotechnology and Biological Sciences Research Council, the Wellcome Trust and the Cancer Research UK. Each time we have written blog posts reporting on questions asked during these discussions, ensuring that the information shared can be used as a future reference for any questions and uncertainties about funders’ policies (Kingsley, 2015a, 2015b, 2016a, 2016b).

## Development of RDM support services

Feedback received from researchers allowed us to start developing services requested by our research community. The key services developed were: a central website with information on RDM, RDM training and support, and a data repository.

### Central website

The first service that we created was a central website on RDM designed to act as one stop shop for all researcher needs: http://www.data.cam.ac.uk. Among many other resources, the website contains online guidance on good data management practice, links to data management support services at Cambridge (including a dedicated data management consultancy appointments and data management plan support service), information on different mechanisms for data sharing and links to discipline-specific data repositories, guidance on funders’ policies, list of FAQs, current training and events on data management and a list of contact points for questions about data management and sharing.

### RDM training and support

Based on needs indicated by our researchers (Johnson et al., 2016), we developed an extensive training offering covering different aspects of RDM and spanning the whole research lifecycle (http://www.data.cam.ac.uk/events). Researchers can have training not only on how to prepare data management plans, how to collect, label and back up their data, but also on how to prepare data for deposit and on how to license research data to ensure maximum re-use. Feedback is collected after each training session to ensure that the modules taught meet researchers’ expectations and needs.

### Data repository

The University of Cambridge established its DSpace institutional research repository, Apollo (http://www.repository.cam.ac.uk), in 2005 (Smith et al., 2003). As a result of feedback received from researchers, a webform was created to allow easy upload of research data. Additionally, since May 2016 each dataset is also assigned a DOI to enable citation and impact measurement. As a result, seven hundred datasets were submitted to the repository since 2015, compared with only 72 data submissions received for a decade from 2005 to 2015 (Teperek, Morgan, Ellefson, & Kingsley, 2016).

Additionally, we also used various communication channels to ensure that researchers are aware of the resources available to them and that our messages are delivered to a wide audience. We took into account the different communication preferences of various stakeholder groups. In addition to having in-person meetings, events and workshops, we also communicated with our academics and support staff via Twitter (https://twitter.com/CamOpenData), newsletters (http://www.data.cam.ac.uk/datanews), e-mails and traditional post.

## Outcomes of the bottom-up approach and lessons learnt

While developing the services to support researchers in RDM, we also changed the way we delivered our information sessions on research data management and sharing. Instead of focusing on funders’ policies and requirements to manage and share research data, we decided to emphasise the personal benefits that could motivate researchers to improve their data management practice and encourage them to share their research data (Markowetz, 2015). Additionally, we organised several events which were researcher-led: instead of administrative staff advocating about the benefits of research data management and sharing, we asked researchers to talk about their own experience with RDM directly to their peers (Teperek, 2016). Researcher-led talks and discussions proved not only to be more compelling to our academic community, but additionally, by inviting researchers who were championing research data management and sharing practice to speak at conferences, we provided them with recognition for their leadership in data sharing.

The uptake of training on RDM exceeded our expectations. Positive feedback received resulted in a growing number of requests for our RDM training support. While the fact that the training delivered was highly valued and met the needs of research community was reassuring, the RDM team consisting of two full-time employees could not meet the growing demand for training across the University. To address this growing demand and also to further recognise and reward researchers who adhere to good data management and sharing practice, a ‘Data Champions’ programme was initiated (Higman, 2016). In this programme, targeted specifically at the research community, researchers were invited to volunteer themselves as research data experts (http://www.data.cam.ac.uk/intro-data-champions). Selected experts were trained by the central RDM team and then became responsible for teaching less experienced colleagues data management skills most relevant to their own research disciplines. The programme not only solves the problem of making discipline-specific training on RDM sustainable, but also helps maintain the engagement within the research community by recognising and rewarding those championing research data management.

Finally, we also focused on designing strategies to maintain the involvement of the broad stakeholder group across the University of Cambridge in RDM service development and delivery. One of the first initiatives here was to ensure that representatives of various communities can formally oversee and contribute to the process of constant improvement of RDM provisions in Cambridge. We launched an open call for people interested in various RDM aspects who would wish to volunteer their time to be part of the RDM Project Group. Encouragingly, over forty applications were received from various stakeholders across the University. Twenty applicants were selected ensuring representation from various departments (archaeology to engineering), different academic (principal investigators, postdocs and students) and non-academic backgrounds (data managers, librarians, research facilitators, administrative and IT officers) and broad expertise areas (information governance, ethics, high performance computing and publishing). The fact that members of the Project Group come from diverse backgrounds not only ensures that the RDM service development is tailored to meet the needs of various stakeholders, but also, through combination of different skillsets and experience of group members, allows constant innovation in our RDM services.

Democratisation of our approaches to RDM had profound effects on the community’s engagement. It not only resulted in an increased number of research datasets submitted to the institutional repository and a growing number of researchers identifying themselves as Data Champions, but also in a change of scope of our discussions with the academic community. Our initial discussions with researchers, which started from debates on whether open data was a waste of time (Teperek, 2015b) shifted to discussions about remaining barriers to sharing (Teperek, 2016) and the benefits of open research (Cadwallader, Jasiewicz, & Teperek, 2016). This suggests that the research community at Cambridge seem to have understood that good data management practice is an integral and necessary part of reproducible research methodology.

## Comparison of the two approaches

In summary, both the top-down and the bottom-up approach have their advantages and disadvantages (Table 1). One of the main advantages of top-down approaches are easy to understand messages, fast service delivery and time-efficiency. On the other hand, top-down approaches are more difficult for the research community to embrace and might lead to community disengagement. Democratic approaches come with numerous benefits (community engagement, trust building), and they are probably the only way to ensure that the services developed are truly aligned with end-user needs. However, one can never underestimate the amount of time and resources required for the successful development and delivery of bottom-up approaches, as well as resources required to maintain the community engagement.

**Table 1.** Comparison of advantages and disadvantages of top-down and bottom-up approaches in service development and delivery.

| Approach | Advantages | Disadvantages |
| --- | --- | --- |
| Top-down, policy-driven approach | Fast service delivery<br>Cost-effective | Risk of community disengagement<br>Risk of solutions misaligned with user needs |
| Bottom-up, researcher-led, democratic approach | Community engagement<br>Services aligned with the user needs<br>Trust between service providers and end users | Time consuming<br>Resource intensive<br>Require careful planning |

The most successful approach in service design and delivery is probably a mixture of top-down and bottom-up approaches. Only by combining the two can one ensure a fast service delivery, while at the same time building a growing base of supportive users. Deciding on an appropriate style of service delivery and approach to communicating with end users should be a key consideration from the beginning of any project developing services. We hope that our lessons learnt might be a useful practical roadmap for other institutions planning to develop or roll-out support services. We believe that these considerations are likely not to be limited to the development and delivery of RDM services in libraries. Our findings and conclusions are likely to be applicable to other institutional services relying on community engagement.

